# Iterative delay correction improves breath-hold cerebrovascular reactivity mapping in clinical populations

**DOI:** 10.64898/2026.04.07.716988

**Authors:** Rebecca G. Clements, Fatemeh Geranmayeh, Niamh V. Parkinson, Molly G. Bright

## Abstract

Cerebrovascular reactivity (CVR), the ability of cerebral blood vessels to dilate or constrict in response to a vasoactive stimulus, is an important measure of cerebrovascular health. Accurate CVR estimation requires accounting for the time required for the vasoactive stimulus to reach each brain region and the time it takes for local arterioles to modulate cerebral blood flow. The temporal search range used to calculate this spatially varying offset can substantially impact CVR estimates, and the appropriate search range may vary across populations, acquisition protocols, and even brain regions. Here, we present an iterative approach for automatically determining the appropriate maximum shift, using breath-hold fMRI data acquired in a cohort of stroke survivors. This approach selectively expands the delay search range only for voxels with estimated delays at the boundary (i.e., near the minimum or maximum shift) until the estimated delay is no longer constrained or a predefined value is reached. In the context of stroke, this approach significantly increased the number of voxels with statistically significant CVR among those initially at the boundary. It also resulted in CVR polarity reversals in voxels originally at the early-response boundary and amplified negative CVR values in voxels originally at the late-response boundary, suggesting that using an iterative maximum shift can critically impact CVR interpretation. This approach is broadly applicable beyond stroke, but careful parameter tuning is required, as illustrated by our demonstration of the parameter tuning process for a participant with Moyamoya disease. Together, these findings suggest that iterative delay correction allows for improved CVR assessments in clinical populations.

## 1. Introduction

Cerebrovascular reactivity (CVR) is the ability of cerebral blood vessels to dilate or constrict in response to a vasoactive stimulus. CVR has numerous clinical applications, including stroke risk prediction (Papassin et al., 2021; Van Niftrik et al., 2024), presurgical planning for brain tumors (Zaca et al., 2011), and prediction of cognitive outcomes (Catchlove et al., 2018; Liu et al., 2024; Sur et al., 2020). One common method for measuring CVR involves using a breath-hold task during blood-oxygen-level-dependent functional MRI (BOLD fMRI) (Bright & Murphy, 2013; Kastrup et al., 2001). Breath holding elevates arterial CO_2_ levels, inducing systemic vasodilation and an increase in cerebral blood flow that can be detected using BOLD fMRI. End-tidal CO_2_ (P_ET_CO_2_), which acts as a surrogate for arterial CO_2_, is measured during the scan, convolved with the canonical hemodynamic response function (HRF), and used as a regressor in a general linear model (GLM) to calculate CVR in each voxel as %BOLD change per CO2 unit change in mmHg.

When calculating CVR, it is important to account for the spatially varying temporal offset between measured P_ET_CO_2_ and the local BOLD response. This offset arises from measurement delays in gas sampling, the time it takes for gases to reach each brain region, and the response times of local arterioles and cerebral blood flow (Stickland et al., 2021). Recent work used a “lagged-GLM” to account for these delays while calculating CVR (Moia et al., 2020; Stickland et al., 2021). This approach involves first correcting for measurement delays and a global vascular transit delay by “bulk-shifting” P_ET_CO_2_ convolved with the HRF to maximize its correlation with the mean time series in gray matter. Then, local hemodynamic delays are accounted for through voxel-wise modeling of the BOLD data using a series of temporally shifted P_ET_CO_2_ regressors, along with nuisance regressors. The voxel-wise P_ET_CO_2_ regressor shift producing the maximum total model R^2^ is considered optimal and used to generate a CVR map. This approach not only improves the accuracy of CVR estimates but also provides complementary information about hemodynamic timing (Stickland et al., 2021).

When using a lagged-GLM to calculate CVR, a temporal lag range must be specified that determines the allowable temporal offsets between P_ET_CO_2_ and the BOLD signal, relative to the bulk shift. In healthy participants, lag ranges of ±9 seconds (s) have been successfully used to calculate CVR (Bailes et al., 2025; Moia et al., 2021; Zvolanek et al., 2025). The choice of lag range for clinical populations is uncertain, as the lagged-GLM has primarily only been validated in healthy participants. Prior studies have reported prolonged hemodynamic delays in cerebrovascular diseases (Donahue et al., 2016; Han et al., 2024; Siegel et al., 2016), suggesting that a larger maximum shift may be appropriate for some populations. However, the applicability of these estimates is unclear, since it is often not clear what the reported delays are relative to or whether they include measurement delays in gas sampling. Additionally, voxel-wise delay estimates derived from different fMRI paradigms (e.g., breath-hold versus resting-state) are not necessarily interchangeable (Gong et al., 2023), suggesting that different shift ranges may be needed for different acquisition strategies. In our preliminary analyses, we observed that simply using a larger maximum shift for an entire dataset can reduce the number of voxels with delay values at the boundaries of the shift range, but it can also result in widespread spurious negative CVR values (Figure 1). This is likely related to the fact that breath-hold tasks can evoke responses resembling a sine wave (Murphy et al., 2011), which are transient in nature and can lead to artifactual negative CVR responses when estimated delays are off by half a period. These findings highlight the importance of carefully selecting the delay search range and suggest that simply using a conservatively large upper bound for all voxels in the brain may not be an appropriate strategy for breath-hold data.

**Figure 1.**
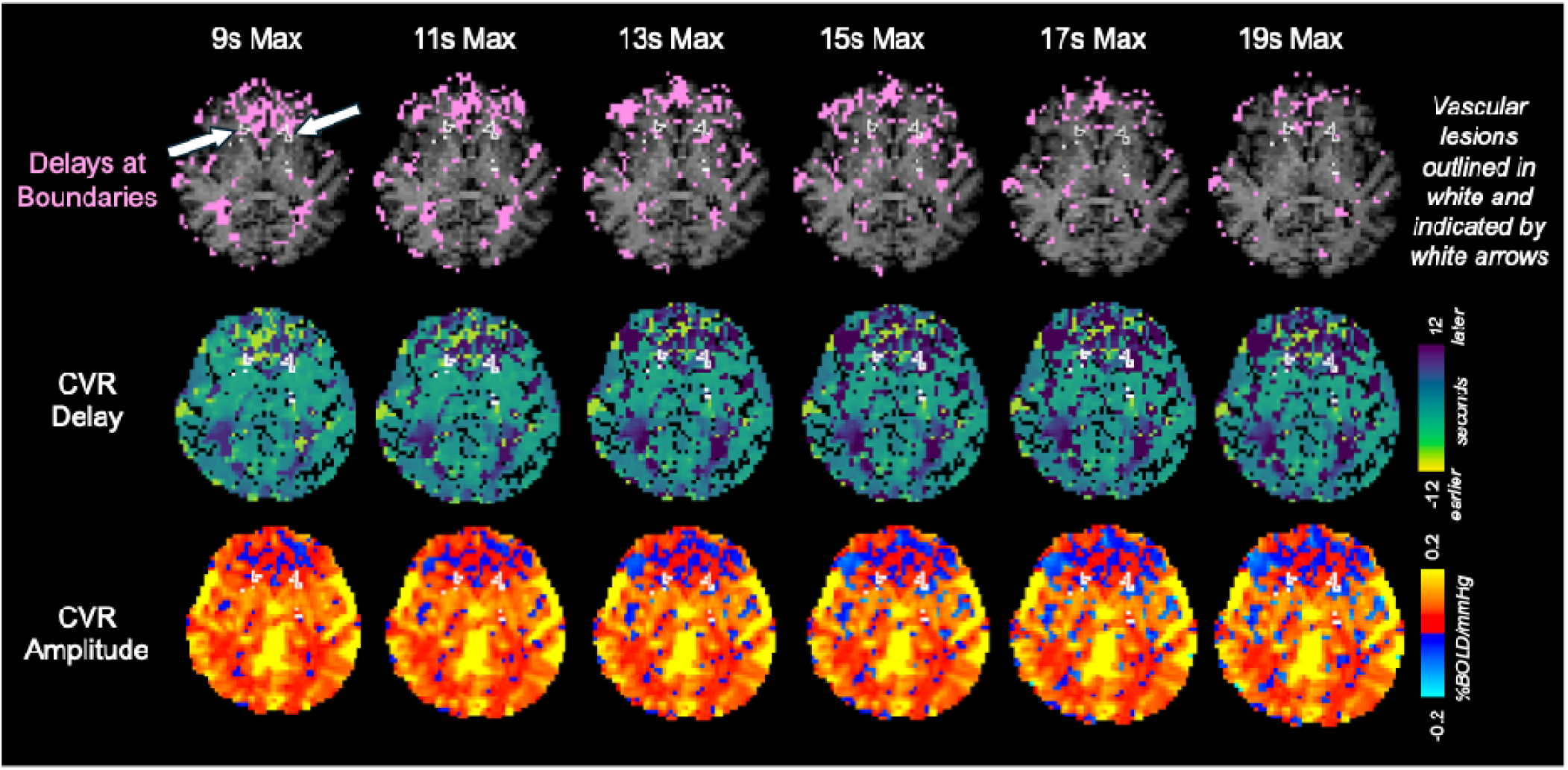
CVR amplitude and delay maps, along with masks of voxels with delay values at the boundaries of the shift range, for one example participant with stroke. A minimum shift of −9 seconds was used, and the maximum shift ranged from 9 to 19 seconds in increments of 2 seconds. We did not explore expanding the minimum shift because delays are calculated relative to normal-appearing tissue, and responses earlier than approximately 9 seconds are not expected in stroke. Vascular lesions are outlined in white and indicated by white arrows. Increasing the maximum shift reduces the number of voxels with delay values at the boundaries of the shift range but also results in more bilateral negative CVR values.

To address these uncertainties, we aim to develop an iterative, voxel-wise approach for determining the appropriate temporal shift range for breath-hold CVR estimation. Estimating the shift range at the voxel level may be particularly advantageous in spatially localized cerebrovascular pathologies, in which hemodynamic delays can vary significantly across brain regions (Braban et al., 2023; Stickland et al., 2021). We propose selectively expanding the maximum shift only in voxels with lag estimates at the boundaries, which may help mitigate the spurious negative CVR values observed in Figure 1. Here, we describe our method and provide open-source code and then validate our method and describe its effects in a cohort of stroke participants.

## 2. Methods

### 2.1. Iterative approach for determination of the appropriate maximum shift

To automatically determine the appropriate lag range, we expanded on the code for *phys2cvr*, an open-source tool for generating CVR maps with a lagged-GLM (Moia et al., 2020, 2024). In addition to the standard parameters used for CVR estimation, the relevant parameters for this analysis are a minimum lag, a starting maximum lag, a final maximum lag, an increment for increasing the maximum lag, and a lag step that determines the resolution of the shifts. These values can be specified by the user and will require careful consideration, depending on the patient population. We suggest that the final maximum lag should be less than half of the breath-hold trial duration, as delays exceeding half of the breath-hold period may introduce spurious negative correlations between P_ET_CO_2_ and the BOLD signal (Moia et al., 2020). For the lag step, previous work has used a lag step that is a quarter of the imaging TR (Stickland et al., 2021).

First, the P_ET_CO_2_ regressor convolved with the HRF, kept at its original sampling frequency without any downsampling, is “bulk shifted” to account for measurement delays and a global vascular transit delay (Moia et al., 2020; Stickland et al., 2021). Bulk shifting is performed by shifting the P_ET_CO_2_ regressor to maximize its cross-correlation with the average time series in normal-appearing gray matter, up-sampled to have the same sampling frequency as P_ET_CO_2_. Next, temporally shifted versions of the bulk shifted P_ET_CO_2_ regressor are generated, spanning lags from the minimum lag to the final maximum lag in increments defined by the lag step, and downsampled to the imaging TR. The BOLD data are then modeled with a separate GLM for each temporally shifted P_ET_CO_2_ regressor, including any nuisance regressors. Using P_ET_CO_2_ regressors with shifts up to the final maximum for this step ensured that the entire range of possible delays was captured, so the lagged-GLM needed to be run only once.

Each voxel’s hemodynamic delay value was estimated by identifying the temporal shift associated with the maximum model fit (R^2^). To identify this maximum, R^2^ values were first examined within the lag window defined by the minimum lag and the starting maximum lag. If the lag corresponding to the maximum R^2^ did not fall within one lag step of either boundary of this window, the lag value for this voxel was considered optimized and no further evaluation of this voxel was performed. If the lag was within 1 lag step of either the positive or negative boundary, the maximum lag was increased by the specified increment and checked again. This process continued until the optimal lag was no longer within one lag step of the boundary or the final maximum lag was reached, whichever came first. An overview of this process for an example subject is shown in Figure 2. After determining the appropriate lag value for each voxel, CVR was calculated as the beta coefficient of the P_ET_CO_2_ regressor convolved with the HRF at its optimal shift. We did not extend the minimum lag because, assuming the bulk shift is estimated from normal-appearing tissue or, in cases of pathology such as Moyamoya disease or carotid stenosis, from unaffected vascular territories, hemodynamic responses more than 9 s earlier than the bulk shift are not expected to be physiologically plausible. Rare exceptions, such as arteriovenous shunting, would require a different analytical approach.

**Figure 2.**
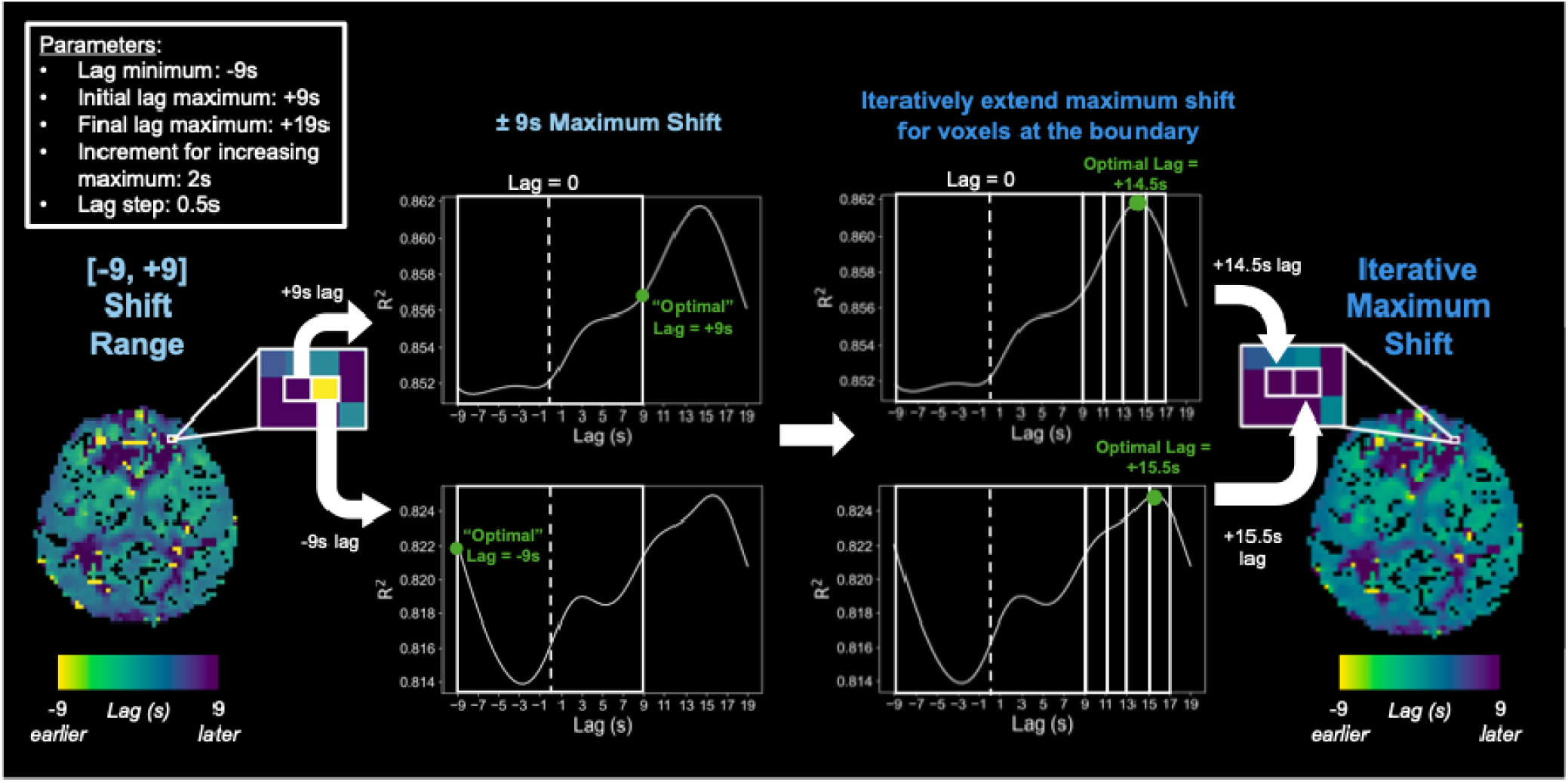
Overview of the iterative delay correction process for 1 example subject, using parameter values shown in the top left. Using a maximum shift of +9 seconds produced “optimal” lag estimates of −9 and +9 seconds in two neighboring voxels. The optimal lag value is identified as the shift producing the maximum R^2^. Because the peak R^2^ occurred at the boundary of the search range, it is likely that the true maximum R^2^ was not captured. For each voxel, the maximum shift was iteratively expanded in 2-second increments until the identified optimal shift no longer occurred at the boundary. This approach yielded lag values of +14.5 and +15.5 seconds in neighboring voxels, which is more plausible since we expect neighboring voxels to have similar delay values.

### 2.2. Validation in a cohort of participants with stroke

#### 2.2.1. Data collection and pre-processing

We validated our methodology using a BOLD fMRI dataset consisting of 64 participants with stroke (49 M, 61 ± 10 years) (mean ± SD) who were imaged at Imperial College London 137 ± 71 days after stroke onset. The participants were recruited as part of the IC3 study (Gruia et al., 2023) approved by the UK’s Health Research Authority (Registered under NCT05885295; IRAS:299333). Informed consent was obtained from all participants for being included in the study. Data collection and processing have been previously described by Clements and colleagues (2026).

Briefly, during a multi-band multi-echo planar imaging sequence (TR = 1.792s, TEs = 13.6c7/40.34/53.71ms, voxel size = 3mm isotropic, MB factor = 2, 154 volumes), participants completed a breath-hold task consisting of 6 repetitions of the following paradigm: 14 s of natural breathing, 16 s of paced breathing, and a 15 s end-expiration breath-hold. Expired CO_2_ was measured during the scan using a sampling frequency of 1000 Hz. T1-weighted whole brain structural image and a fluid-attenuated inversion recovery (FLAIR) image were also acquired. As previously described (Clements et al., 2025; Zvolanek et al., 2023), 28 participants were excluded due to having low-quality P_ET_CO_2_ timeseries, making the final sample size 36 participants (28 M, 10 participants with hemorrhagic stroke, 63 ± 12 years). It is important to note that this level of attrition is conservatively high. This approach would likely still be effective using surrogate measures of P_ET_CO_2_, like respiration volume per time (Zvolanek et al., 2023) or deep-learning derived P_ET_CO_2_ (Clements et al., 2025), but we wanted to focus on and validate this approach in participants with high-quality measured P_ET_CO_2_ first.

Pre-processing of the BOLD fMRI data included volume registration, distortion correction, brain extraction, optimal combination of the 4 echoes with *tedana* (DuPre et al., 2021; Kundu et al., 2012, 2013), and smoothing with a 4mm FWHM kernel (Cox, 1996; Jenkinson et al., 2012). Multi-echo independent component analysis (ME-ICA) was performed with *tedana* to produce additional nuisance regressors, using methods described by Clements and colleagues (2026). The expired CO_2_ data were processed by using in-house MATLAB code to identify end-tidal peaks, manually verifying the peaks, and linearly interpolating the peaks to create P_ET_CO_2_ timeseries with the same sampling frequency as the original CO_2_ data. P_ET_CO_2_ timeseries were then convolved with canonical HRF (Friston et al., 1998).

#### 2.2.2. CVR calculation and thresholding

CVR maps were generated using our modified version of *phys2cvr* that iteratively determines the appropriate lag range. The pre-processed BOLD data were modeled using temporally shifted variants of P_ET_CO_2_ convolved with the HRF, motion parameters and their derivatives (obtained during volume registration), Legendre polynomials up to 3^rd^ order, and ME-ICA components classified as noise that were orthogonalized to the ME-ICA components classified as signal and all other regressors in the model. Orthogonalization of the ME-ICA components was performed in alignment with the previously validated “conservative” ME-ICA approach developed by Moia and colleagues (2021) for CVR analyses. The lag minimum and starting lag maximum were set to [-9, +9] s and the lag step was set to 0.5 s, consistent with values used previously in healthy participants (Bailes et al., 2025; Moia et al., 2021; Zvolanek et al., 2025). The final lag maximum as set to 19 s and the increment for increasing the lag maximum was set to 2 s. For comparison, CVR maps were also generated using the standard *phys2cvr* approach with a fixed lag range of [-9, +9] s and a lag step of 0.5 s.

To characterize changes in CVR for voxels initially at the delay boundary using a [-9, +9] s lag range, we identified voxels whose lag values were within one lag step (0.5 s) of the minimum or maximum shift. Specifically, voxels were classified as being at the early response (−9 s) boundary if their lag was less than or equal to −8.5 s and at the late response (+9 s) boundary if their lag was greater than or equal to +8.5 s. We also identified voxels with statistically significant CVR. CVR values were considered significant if they had absolute *t*-statistics greater than 1.98 (corresponding to *p*<0.05). This value was further adjusted using Šidák correction to account for running 57 different models for the iterative maximum approach ([-9,19] s search range in 0.5 s increments) and 37 different models for the standard approach ([-9,9] s search range in 0.5 s increments) (Bright et al., 2017; Šidák, 1967). We note that this threshold is likely conservative for the iterative maximum approach, since we did not use a lag maximum of +19 s for all voxels.

#### 2.2.3. Generation of region of interest masks

To evaluate the effects of our iterative maximum approach across different tissue types, we generated region of interest (ROI) masks using FSL tools (Jenkinson et al., 2012). Stroke lesions and white matter hyperintensities, which we will refer to collectively as vascular lesions, were manually delineated using the FLAIR images and verified by neurologist FG. The T1-weighted image was brain-extracted with *BET* and segmented into gray matter (GM), white matter (WM), and cerebrospinal fluid (CSF) with *FAST* using a threshold of 0.5 (Y. Zhang et al., 2001). Vascular lesion masks were transformed to T1-space using *FLIRT* (Jenkinson et al., 2002; Jenkinson & Smith, 2001) and subtracted from the GM, WM, and CSF masks to generate masks of normal-appearing GM, WM, and CSF, which were then transformed to functional space.

Because negative CVR values have been observed in and around the ventricles, and prior work suggests that CVR mechanisms in these regions may differ (Bright et al., 2014; Thomas, Liu, Aslan, et al., 2013), we wanted to evaluate the ventricles and peri-ventricular space separately from the remaining CSF. For each subject, we transformed a mask of the ventricles from the N27 template brain (Holmes et al., 1998) to functional space. This mask was dilated by 1 level using AFNI’s *3dmask_tool* (Cox, 1996) and the original functional mask of the ventricles was subtracted to generate a mask of periventricular space. To avoid having overlapping voxels in the ROI masks, voxels in the ventricles or in periventricular space were excluded from the GM, WM, and CSF masks, and vascular lesion voxels were excluded from the masks of the ventricles and periventricular space. This analysis resulted in 5 total ROI masks in functional space: normal-appearing GM, normal-appearing WM, CSF, ventricles, peri-ventricular space, and vascular lesions.

### 2.3. Example of parameter tuning for a different pathology

To assess how the iterative delay correction approach generalizes beyond stroke, and to illustrate how parameters may need to be adjusted for other cerebrovascular pathologies, we applied our methods to data from a 31-year-old male with unilateral Moyamoya disease causing an occluded right middle cerebral artery (MCA). This dataset was collected at Northwestern University under a study approved by the Northwestern University Institutional Review Board. Informed consent was obtained from the participant for being included in the study. The participant completed a breath-hold task during a multi-band multi-echo gradient-echo echo planar imaging sequence (TR = 1.5s, TEs = 10.6/27.83/45.06/62.29/79.52 ms, voxel size = 2.5mm isotropic, MB factor = 4, 212 volumes). The breath-hold task consisted of 6 repetitions of the following paradigm: 24 s of paced breathing, a 15 s hold, a 2 s exhale, and 8 s of recovery. Afterwards, the participant completed another 24 s of paced breathing. During the scan, expired CO_2_ was measured using a nasal cannula and used to calculate end-tidal CO_2_. The breath-hold task and fMRI pre-processing steps were similar to those used for the stroke dataset.

For this participant with right MCA occlusion, we first evaluated the reference signal used to calculate the bulk shift. The bulk shift is the initial realignment applied to the P_ET_CO_2_ regressor to maximize its correlation to the reference signal. It accounts for measurement delays and a global transit delay and is applied before calculating voxel-wise hemodynamic delays. We tested two different ROIs for generating the reference timeseries: a mask of all normal-appearing gray matter and a mask of normal-appearing gray matter excluding the affected vascular territory. The affected vascular territory was identified using a mask of the right MCA from a vascular territory atlas (Schirmer et al., 2019a, 2019b) and transformed into functional space. The main benefit of the whole-brain gray matter mask is that it can be more readily obtained and does not require *a priori* knowledge of pathology. At the same time, it also may generate a reference timeseries that is temporally “blurred” compared to the gray matter mask that excluded the affected vascular territory, as it includes voxels with a large range of delay values (Tong et al., 2019). Compared to the gray matter mask that excluded the affected vascular territory, the timeseries in the whole-brain gray matter mask could also be reduced in amplitude due to averaging across regions with reduced or negative responses to the breath holds.

The increment for increasing the maximum lag was set to 2 s, equivalent to our TR rounded to the nearest whole number. Because we did not expect any voxels to exhibit responses more than 9 s earlier than the unaffected vascular territories, the minimum lag value was set to −9 s. The final maximum lag value should be less than half of the breath-hold trial length (49 s), and thus was set to 23 s. Finally, to determine the appropriate starting maximum, we experimented with using starting maximum values of 9, 11, 13, 15, 17, and 19 s.

## 3. Results

### 3.1. Validation in a cohort of participants with stroke

When using a ±9 s shift range, the median proportion of voxels at the −9 and +9 s boundaries was 4% and 5% of the brain, respectively, with some participants showing up to 18% at −9 s and 13% at +9 s. (Figure 3A). Most of these voxels were nonsignificant. Using an iterative maximum shift, voxels at the late response boundary mostly shifted to slightly later delays, with only a minority (25%) remaining at the original boundary (Figure 3B). The largest proportion of voxels had an appropriate maximum delay of +11 s, and progressively fewer voxels required longer delays than 11 s. Most voxels at the early response boundary (64%) retained their original early delay, while the remainder shifted from the early boundary to later delays. Using an iterative maximum significantly increased the percentage of significant voxels among voxels initially at the late response boundary and the early response boundary, as indicated using a 1-sided paired *t*-test (*p*<0.05, Bonferroni corrected). Importantly, these significant increases occurred despite applying a stricter Šidák correction for the iterative maximum approach. This result suggests that using an iterative maximum can recover significant CVR estimates for voxels with prolonged delays.

**Figure 3.**
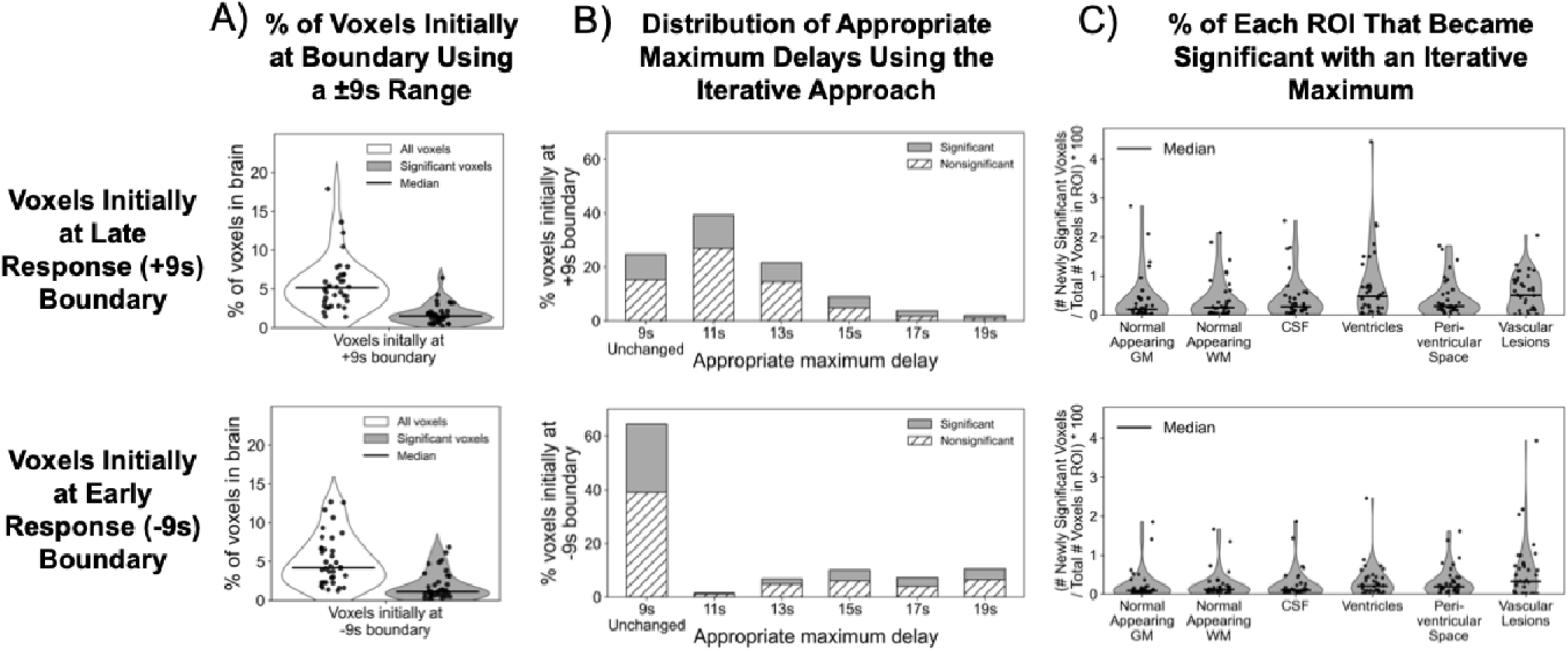
Effects of the iterative approach for voxels initially at the lag boundaries with a [-9, +9] shift range. The top row describes voxels initially at the late (+9 seconds) response boundary, and the bottom row describes voxels initially at the early (−9 seconds) response boundary. (A) shows the percentage of voxels in the brain that were initially located at each lag boundary across subjects. Significant voxels were identified using a threshold of *p*<0.05 and Šidák correction. (B) shows the distribution of the appropriate maximum delay identified by the iterative maximum approach for voxels initially at the lag boundaries, across all the subjects. (C) shows the percentage of voxels in each ROI that became significant with an iterative maximum.

We also investigated the location of voxels that switched from being nonsignificant using a +9 s maximum shift to significant using an iterative maximum shift, within the predefined ROI masks (Figure 3C). For both voxels initially at the late response boundary and voxels initially at the early response boundary, the percentages of voxels that became significant were relatively similar across ROIs. However, vascular lesions, which included both lesions and white matter hyperintensities, showed the highest percentage. This is consistent with the expectation that stroke lesions will exhibit longer hemodynamic delays (Braban et al., 2023). White matter hyperintensities have also been previously associated delayed arterial transit times (R. Zhang et al., 2022) and thus may also exhibit prolonged hemodynamic delays.

Next, we evaluated how the iterative maximum approach impacted CVR amplitude values (Figure 4). For this analysis, we only included voxels that were statistically significant using delay correction with an iterative maximum shift. For voxels that were initially at the late response boundary, negative CVR values became slightly more negative, and values near zero shifted toward more negative CVR amplitudes (Figure 4A). This effect is illustrated in example CVR maps (Figure 5, Example 1) which show a brain region where, with the iterative approach, CVR becomes more negative and statistically significant, and the estimated hemodynamic delay is increased. Negative CVR values may represent effects such as vascular steal (Poublanc et al., 2013; Venkatraghavan et al., 2018), oxygen metabolism upregulation (Arteaga et al., 2014), or fractional changes in fluid volume (Blockley et al., 2011; Bright et al., 2014; Thomas, Liu, Aslan, et al., 2013).

**Figure 4.**
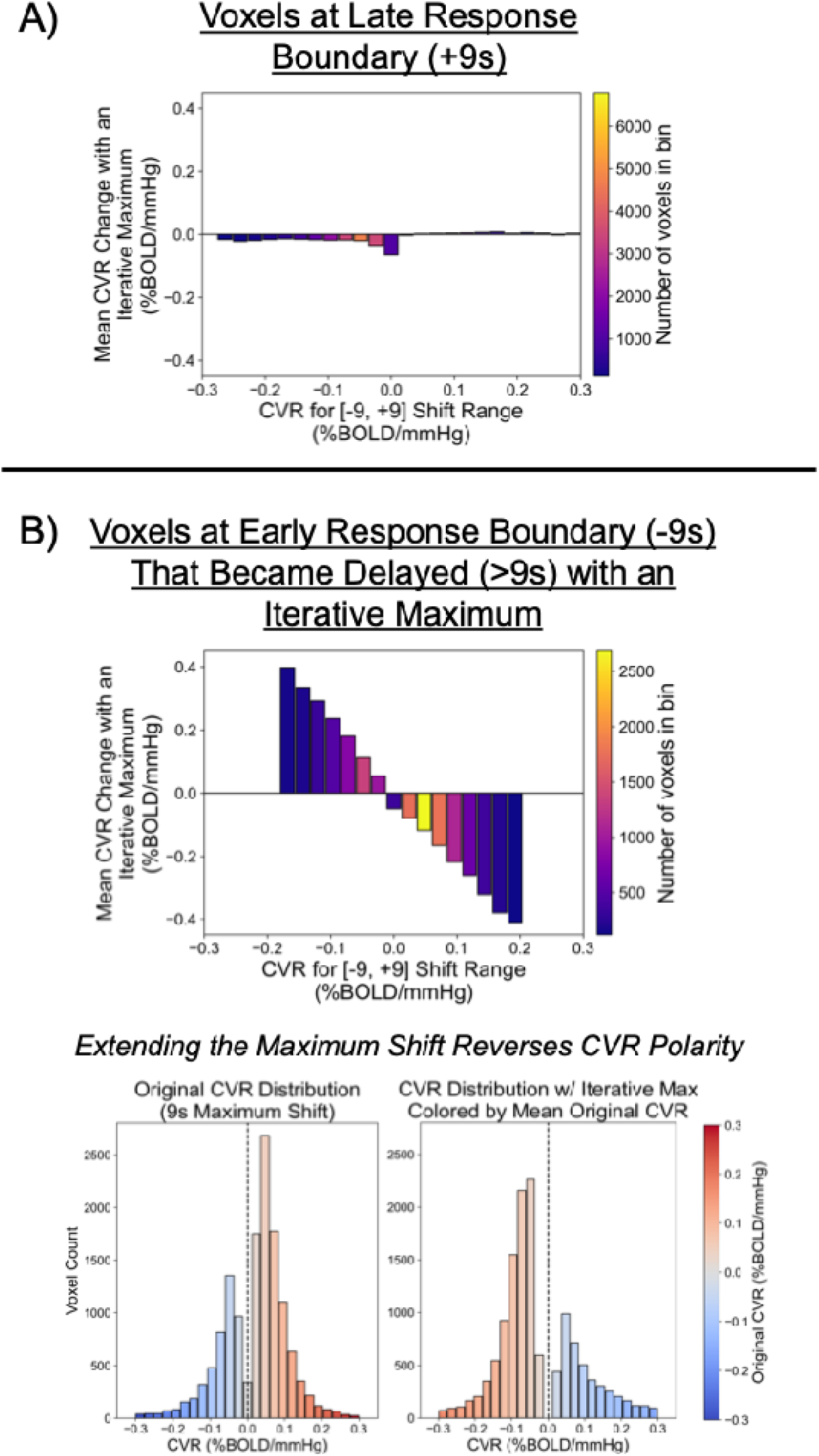
Changes in CVR amplitude with an iterative maximum shift for voxels at the late response boundary (A) and the early response boundary that became delayed with an iterative maximum (B). For the top two plots, CVR values were divided into 25 bins ranging from −0.3 to 0.3, and the average change in CVR across subjects was calculated for each bin. Bins containing fewer than 100 voxels were excluded. Voxels at the late response boundary with negative or near-0 CVR values tended to have CVR values that became slightly more negative. Voxels at the early response boundary that switched to being delayed with an iterative maximum reversed in CVR polarity. This effect is illustrated in the bottom two plots: the left plot shows the original CVR distribution with a 9 s maximum shift, and the right plot shows the CVR distribution after the iterative maximum shift. Voxels are colored according to their original CVR values, and the color reversal between the two plots demonstrates the polarity change.

**Figure 5.**
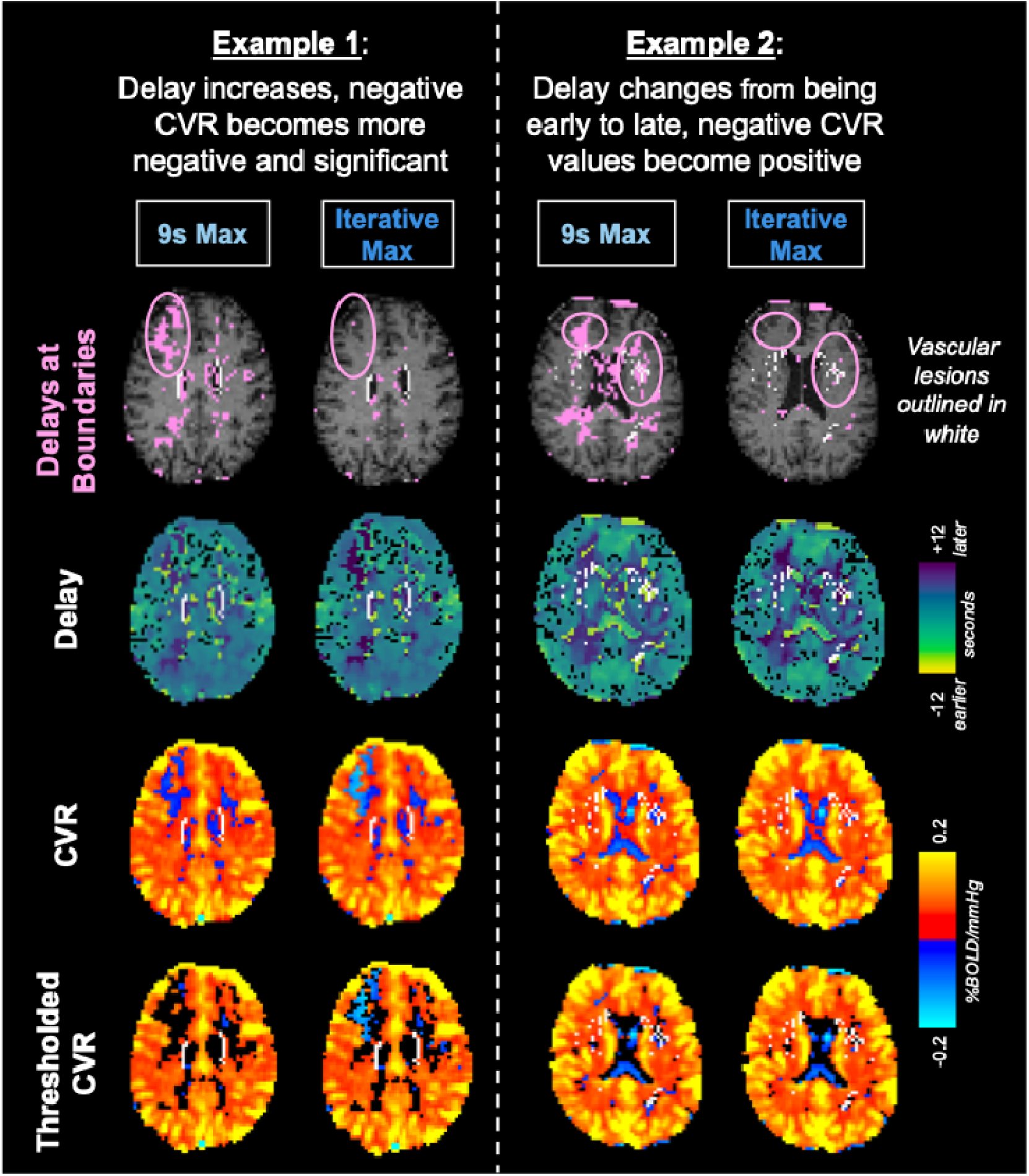
Example CVR maps for 2 example subjects, generated using a 9 second maximum shift and an iterative maximum shift. The top row shows voxels with delay values at the boundaries of the tested lag range (pink). The pink circle indicates brain regions of interest, where the text description at the top applies. The remaining rows show delay maps, CVR amplitude maps, and thresholded CVR maps, generated using a threshold of *p*<0.05 and Šidák correction.

For voxels that were initially at the early response boundary that became delayed (>9s delay) with an iterative maximum, we observed relatively large positive changes for CVR values that were initially negative, and large negative changes for CVR values that were initially positive (Figure 4B). To determine whether this indicated a reversal in CVR polarity, we plotted the distribution of CVR values obtained with the 9 s maximum delay and the distribution obtained with the iterative maximum, colored by the original 9s delay CVR values (Figure 4B, bottom). The inversion of colors in the distribution plots indicates that extending the delay range for voxels at the early-response boundary often resulted in flipped CVR polarity. This effect is also evident in an example CVR map (Figure 5, Example 2), in which, using an iterative maximum, negative CVR values became positive and CVR delay changed from being early to late. In Example 2, the voxels that changed CVR polarity and switched from having early to late responses were near vascular lesions, in line with previous studies reporting longer delays in these regions (Geranmayeh et al., 2015).

These findings suggest that the iterative delay correction approach can critically impact CVR interpretation. Because delays were measured relative to normal-appearing tissue, we did not extend the early-response boundary because we did not expect any voxels to respond more than 9 s before normal-appearing tissue. Future work should validate our findings using controlled gas delivery with longer hypercapnic periods, where delay correction is expected to be less critical. It would be informative to determine whether regions that switched from very early to very late responses using an iterative maximum in breath-hold data also had delayed responses under sustained hypercapnia. Additionally, future validation using perfusion-weighted MRI measurements would help establish that delays measured using the iterative approach more accurately reflect underlying hemodynamics.

### 3.2. Example of parameter tuning for a different pathology

In the participant with unilateral Moyamoya disease, we calculated the bulk shift using a mask of all normal-appearing gray matter and a similar mask that excludes the affected vascular territory, and we found that these masks produced nearly the same results (bulk shifts of 17.1 and 17.3 s, respectively). We decided to use the mask of gray matter in the unaffected vascular territories, only, since it is expected to be slightly more accurate. However, these observations suggest that the bulk shift calculation in our lagged-GLM is sufficiently robust to widespread variations in hemodynamic delay and we may not require *a priori* knowledge of cerebrovascular pathology.

After the bulk shift is applied to account for measurement delays and a global transit delay, voxel-wise hemodynamic delays are determined using the specified shift range, Using a minimum and final maximum lag of −9 and +23 s, respectively, and increasing the maximum lag in 2 second increments, we experimented with different starting maximum values (Figure 6). In the affected vascular territory, outlined in white, we observed that using a starting maximum of 9, 11, or 13 s resulted in a region of white matter having shorter delays (around 0-5s) than nearby gray matter (around 9-14s) (see white arrows). Although the validity of this result is uncertain, white matter typically has a slower CVR response than gray matter (Thomas, Liu, Park, et al., 2013), and Moyamoya is anticipated to specifically lengthen vascular transit times, suggesting that we have an insufficient initial search range to capture all voxels with longer hemodynamic delays. Using a starting maximum of at least 15 s resulted in more appropriate (i.e., longer) estimated delays in this specific white matter area relative to gray matter across the affected vascular territory, which is more plausible.

**Figure 6.**
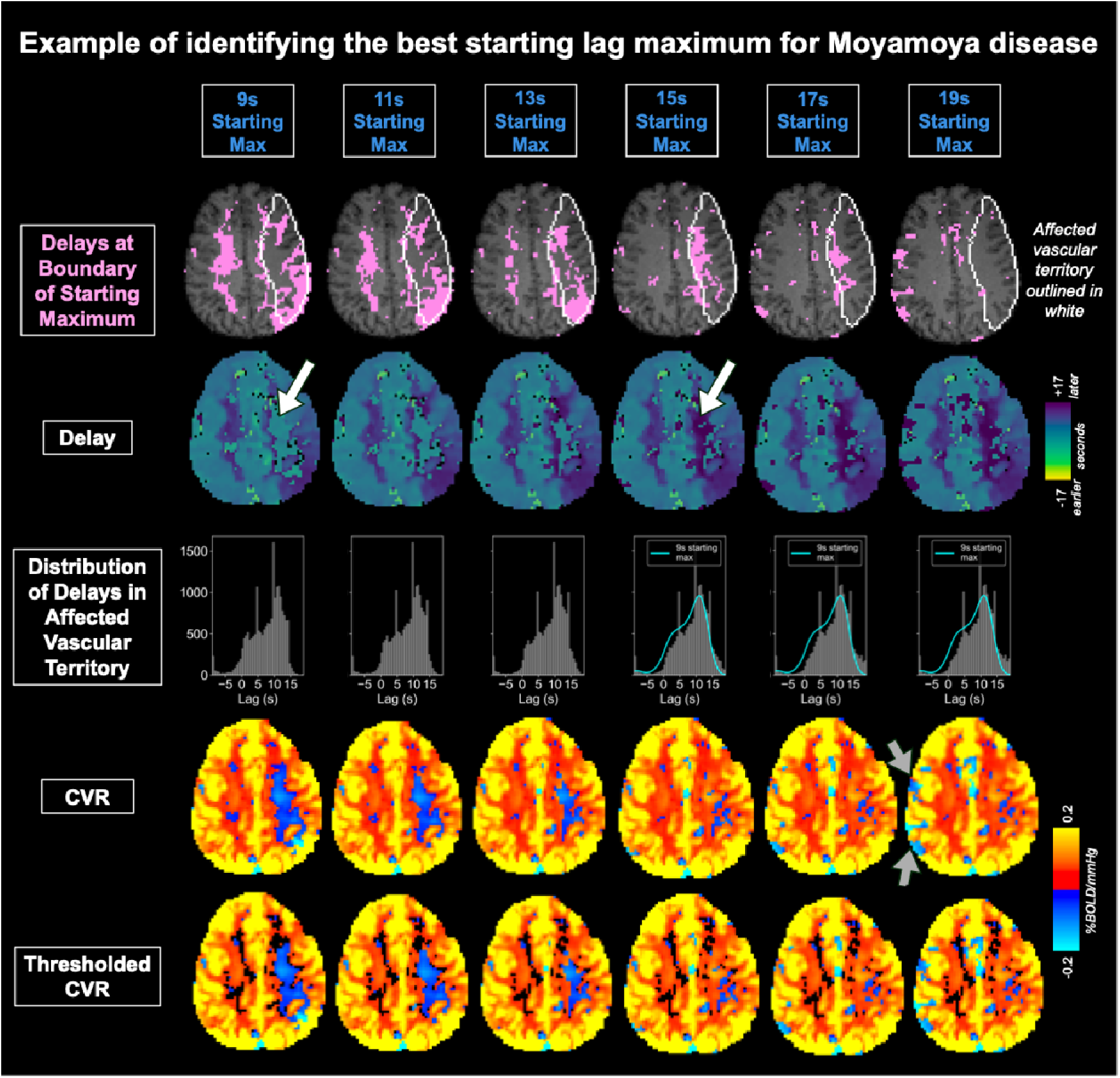
CVR maps generated using an iterative maximum for a participant with unilateral Moyamoya disease. Columns correspond to different starting maximum delays (9-19 seconds, in 2 second increments). As the starting maximum increases, delays within the affected vascular territory become longer and more spatially consistent, particularly in white matter (see white arrows in row 2). Histograms of delay values within the affected territory demonstrate that the peak at 0–5 seconds observed with a 9-second starting maximum diminishes with higher starting maxima and is replaced with more delay values between 10–15 seconds. A starting maximum of 19 s resulted in more negative CVR values outside of the affected vascular territory (see gray arrows), suggesting that a 15 or 17 s starting maximum may be most appropriate. Thresholded CVR maps were generated using a threshold of *p*<0.05 and Šidák correction.

This increase in delays in white matter is also illustrated by the distribution of delays in the affected vascular territory (see row 3 of Figure 6). Specifically, the small peak between 0-5 s observed with a 9 s starting maximum disappears when using an initial maximum shift of 15 s or 17 s and is replaced with more delays in the 10-15 s range. This shift is evident when comparing the envelope of the 9 s delay histogram with the corresponding histograms for the 15, 17, and 19 s starting maxima (see histograms in columns 4, 5, and 6 in Figure 6). These findings suggest that when the starting maximum is too short, false negative CVR estimates may occur because the estimated delay is misaligned by approximately half of a breath-hold period. In these cases, the selected delay may not be at the boundary, so the search range would not be expanded despite the incorrect alignment. We also observed that using a starting maximum of 19 s resulted in more negative CVR values in brain areas outside of the affected vascular territory (see gray arrows), suggesting that a 19 s starting maximum also may not be appropriate, akin to our results in stroke at a 15 s starting maximum (Fig. 1). These findings suggest that using a starting maximum of 15 or 17 s produce optimal results and may be best for this pathology.

In summary, these results demonstrate that while the iterative delay correction is designed to be applied to any cerebrovascular pathology, careful consideration of parameter values is essential. For participants with Moyamoya disease, a larger starting maximum appears necessary to accurately capture hemodynamic delays within the affected vascular territory. This likely reflects the more widespread and severe delays characteristic of Moyamoya disease, which involve an entire vascular territory, in contrast to the more localized effects we observed in stroke. In the future, leaving more time between breath-hold trials (potentially around 1 minute, though the optimal duration may vary) may also be helpful for patient populations with longer delays and may make successful delay correction less dependent on specific parameter values. Future work should involve more extensive validation in larger datasets to more confidently determine optimal parameter values for iterative delay correction for populations of interest.

## 4. Conclusion

Accounting for voxel-wise hemodynamic delays is critical for accurate CVR estimation, particularly in clinical populations with prolonged vascular responses. The choice of maximum temporal shift can greatly impact calculated hemodynamic delay values and may need to vary across populations and acquisition protocols. We presented an approach for determining the appropriate maximum delay value that involves iteratively extending the delay range only for voxels with lag values at the boundary of the initial search window for the optimal lag. In participants with stroke, this approach significantly increased the number of voxels with statistically significant CVR among those initially at the boundary. For voxels initially at the late-response boundary, voxels with 0 or negative CVR values became slightly more negative, increasing confidence that these values reflect physiological signals rather than noise. In contrast, for voxels at the early response boundary, an iterative maximum resulted in a reversal of CVR polarity, greatly altering interpretation of these responses.

Importantly, our approach can be readily applied across different populations, although our choice of parameter values has only been validated in stroke. Careful consideration of parameter values, such as the starting and final maximum lag and the reference timeseries for calculating the bulk shift, is necessary when moving to other cohorts. To guide future work, we demonstrated the parameter tuning process for unilateral Moyamoya disease and showed that certain parameter values may need to be altered to account for more extreme or widespread hemodynamic delays. Together, the findings of this study highlight the importance of appropriately accounting for hemodynamic delays in CVR analysis and demonstrate that an iterative, data-driven approach for determining the delay search range can improve the accuracy and interpretation of CVR estimates.

## Data & Code Availability

Code for calculating CVR using the iterative delay correction approach is available at https://github.com/BrightLab-ANVIL/phys2cvr_Iterative and is being submitted as a pull request on GitHub to be incorporated into the main phys2cvr code (https://github.com/smoia/phys2cvr). Imaging data are not publicly available as they are part of a study that is still ongoing. Access may be considered subject to inter-institutional agreements once the study objectives have been achieved.

## Author Contributions

Rebecca G. Clements: Conceptualization, Methodology, Software, Formal analysis, Investigation, Data curation, Writing—original draft, Writing—reviewing and editing, Visualization, Project administration. Fatemeh Geranmayeh: Methodology, Resources, Data Curation, Writing—reviewing and editing, Supervision, Funding acquisition. Niamh V. Parkinson: Investigation, Project administration. Molly G. Bright: Conceptualization, Methodology, Resources, Writing—review & editing, Supervision, Project administration.

## Acknowledgements

We are grateful to all the participants for their time and involvement in the study. We thank all the research assistants and students for their help with data collection and manual stroke lesion segmentation.

